# Characterizing JP-5 Biodegradation Potential in Hawaii Soil Microbiome: Phylogeny, Growth Kinetics, and Biosurfactant Production

**DOI:** 10.1101/2025.11.21.689844

**Authors:** Yangyang Zou, Min Ki Jeon, Tao Yan

**Affiliations:** Water Resources Research Center, University of Hawaii at Manoa, 2540 Dole Street, Holmes Hall 283, Honolulu, HI 96822, USA; Department of Civil, Environmental and Construction Engineering, University of Hawaii at Mānoa, 2540 Dole Streets, Holmes Hall 383, Honolulu, HI 96822, USA

**Keywords:** JP-5 biodegradation, Hydrocarbon degrading microorganisms, soil microbiome, growth kinetics, Biosurfactant production, 16S rRNA phylogenetic

## Abstract

Soil and groundwater contamination from petroleum spills continues to present major environmental and human health challenges. A recent fuel contamination event in the Pearl Harbor region of the Island of Oahu, Hawaii, highlights the need for a better understanding of the biodegradation capabilities of the indigenous soil microbiome and potential for bioremediation. Soil sampling and subsequent cultivation-based enumeration showed that the abundance and relative abundance of JP-5 degrading bacterial biomass to be 1.9 × 10^4^ MPN/g (σ=3.4 × 10^4^ MPN/g) and 2.4 % (σ= 4.4 %), respectively. Isolation and 16S rRNA gene sequencing showed diverse phylogenetic groups of JP-5 degrading bacteria, including *Achromobacter*, *Ralstonia*, *Stenotrophomonas*, *Pseudomonas*, *Acinetobacter*, *Staphylococcus*, *Bacillus*, and *Gordonia*. Logistic modeling of batch growth experiments revealed significant variation in growth kinetic parameters amongst the soil bacterial isolates. Biosurfactant production by the JP-5-degrading bacterial isolates showed significant correlation with maximum optical density (*r* = 0.69, *p* < 0.001), relative growth rate (*r* = 0.41, *p* = 0.008), and relative lag phase duration (*r* = –0.36, *p* = 0.02), indicating an important role of biosurfactant production in enhancing hydrocarbon bioavailability. The results revealed the ubiquitous presence and phylogenetic diversity of JP-5 degrading microbial capabilities in the Hawaii soil microbiome and their potentials for bioremediation applications.

## Introduction

Leakage of raw petroleum and refined petroleum products during storage and transportation continues to pose a significant threat to environmental and ecological health. A notable incident occurred in 2021 in the Pearl Harbor region of the Island of Oahu, where a large amount of jet fuel (JP-5) was accidentally released from underground storage tanks (USTs), contaminating the surrounding soil and entering the underlying groundwater aquifer. The leaked fuel contains diverse petroleum hydrocarbon compounds (PHCs), including paraffins, cycloparaffins, aromatics, and olefins (Jung et al., 2002). Direct exposure to these PHCs has been associated with adverse effects on the respiratory, immune, and nervous systems (Smith et al., 2010), while indirect health risks may include the formation of disinfection byproducts when the PHC-contaminated water is subjected to chlorination during drinking water treatment (Brinkmann et al., 2024).

Because of the observed persistence of PHCs at other sites (Essaid et al., 2011), long-term stewardship of the contamination site is required, and it is imperative to explore potential remediation options in order to safeguard the sole-source groundwater aquifer of the region. Traditional remediation technologies, including physical and chemical methods, are often associated with high operational costs, with the potential to introduce secondary pollution. On the other hand, bioremediation offers a cost-effective and environmentally sustainable alternative (Pandolfo et al., 2023). Studies at other petroleum contamination sites have reported diverse populations of indigenous soil microbiomes with the capabilities to efficiently degrade petroleum hydrocarbons, often utilizing them as carbon and energy sources (Chunyan et al., 2023; Li et al., 2022; Meintanis et al., 2006; Mohammed et al., 2023), including *Pseudomonas* (Medic et al., 2020; Posada-Baquero et al., 2020; Varjani & Upasani, 2016), *Achromobacter* (Chunyan et al., 2023), *Acinetobacter* (J. Czarny et al., 2020; Mohammed et al., 2023; Pandolfo et al., 2023), *Bacillus* (Li et al., 2022; Liu et al., 2016; Wang et al., 2019), and *Rhodococcus* (Diallo et al., 2021; Song et al., 2011; Van Hamme et al., 2003).

Different indigenous soil bacterial species can exhibit different target hydrocarbon substrates and degradation efficiencies. For example, *Achromobacter* species has been shown to degrade C12-C28 alkane with an efficiency of 78% within 7 days (Chunyan et al., 2023), while *Bacillus* can degrade approximately 53% of C13-C30 alkane over 28 days (Li et al., 2013). Some soil bacteria can degrade more recalcitrant components of PHCs; e.g., some *Pseudomonas* species have demonstrated the ability to degrade polycyclic aromatic hydrocarbons (PAHs) such as fluorene, phenanthrene, and pyrene within 7 days with degradation efficiency of 96%, 50%, and 40% respectively (Medic et al., 2020). However, few studies have examined the microbiomes of tropical Hawaiian soils exposed to jet fuel contamination, and it remains uncertain whether results from temperate regions can be directly applied to these distinct, volcanic, and highly weathered soil environments.

This study aims to characterize the microbial potential for JP-5 jet fuel biodegradation in the surface soil on the Island of Oahu. The abundance of JP-5-degrading bacteria in soil samples collected from Oahu Island, Hawaii, was determined to provide baseline information on the natural occurrence of hydrocarbon-degrading microbial populations. The phylogenetic affiliation of the JP-5 degrading bacterial isolates were determined through 16S rRNA gene sequencing. The growth kinetics of the bacterial isolates were experimentally determined in batch growth experiments and were modeled by using the logistic growth model to determine key parameters, including maximum optical density, relative growth rate, and relative lag phase duration. Lastly, the biosurfactant production potential of the soil bacteria was quantified and then compared with their growth kinetics with JP-5 as the sole carbon source and their phylogenetic affiliation.

## Materials and Methods

### Soil sampling and characterization

Soil samples were collected from 12 sites, one forest site (F) and one urban site (U) from six watersheds at the southern shore of the Island of Oahu, including Pālolo (PL), Mānoa (MN), Pauoa (PU), Nu uanu (NU), Kalihi (KL), and Kapālama (KP) (**Figure 1**). The sample IDs were designated as combinations of sampling site and soil type (e.g., Pālolo forest: PLF). Soil samples were collected from the surface layer (0–20 cm depth) at locations approximately 20 cm to 100 cm from the stream edge; GPS coordinates for each site are provided in **Table S1**.

**Figure 1.**
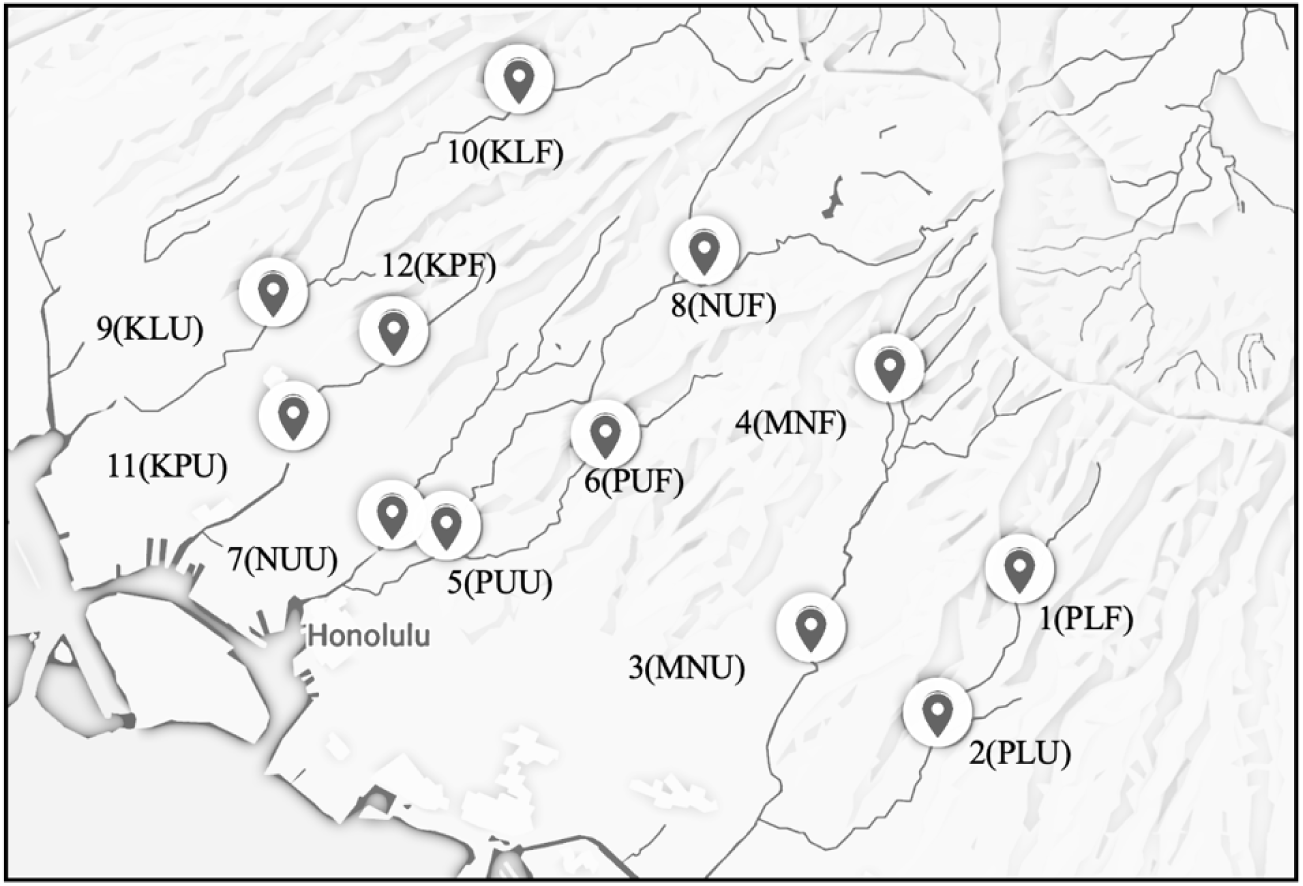
Soil sampling locations at urban and forest sites at six watersheds on the Island of Oahu, Hawaii.

### Bacterial enumeration

The abundance of the total bacterial population and the JP-5-degrading bacterial sub-populations were determined using a most probable number (MPN) approach (Papen & Von Berg, 1998). To estimate total bacterial abundance, 1 g of the soil sample was inoculated into 9 ml Reasoner’s 2A (R2A) medium (composition: yeast extract 0.5 g/L, proteose peptone No.3 0.5 g/L, casamino acids 0.5 g/L, glucose 0.5 g/L, soluble starch 0.5 g/L, K_2_HPO_4_ 0.3 g/L, MgSO_4_·7H_2_O 0.05 g/L, sodium pyruvate 0.3 g/L) (Gibbs & Hayes, 1988), and vortexed for 2 min to obtain a 10^-1^ stock suspension, followed by serial 10-fold dilutions up to 10^-9^. Dilution factors refer to tenfold serial dilutions of the soil suspension used for MPN enumeration. To quantify JP-5-degrading bacteria, 1 g of soil sample was inoculated into 9 mL of sterile Bushnell-Haas (BH) medium (composition: MgSO_4_ 0.2 g/L, CaCl_2_ 0.02 g/L, KH_2_PO_4_ 1 g/L, K_2_HPO_4_ 1 g/L, (NH_4_)_2_SO_4_ 1 g/L, FeCl_3_ 0.05 g/L) (Bushnell & Haas, 1941), followed by serial dilutions from 10^-1^ to 10^-6^, and supplemented with 1% (v/v) JP-5 jet fuel as the sole carbon source. Dilution factors refer to tenfold serial dilutions of the soil suspension used for MPN enumeration.

The R2A MPN tubes were incubated at 28°C, and optical density at 600 nm wavelength (OD_600_) was measured at 48, 72, and 168 hours using a spectrophotometer (Eppendorf, Hamburg, Germany) (Reasoner & Geldreich, 1985). The BH MPN tubes were incubated at 30°C with shaking at 180 rpm for up to 4 weeks (Araújo et al., 2020), with OD_600_ measurements every week. A MPN tube was considered positive for bacterial growth if it satisfied at least two of the following criteria: (1) a statistically significant increase in OD_600_ (*p* < 0.05, *t*-test) from the preceding sampling point, (2) a change in culture medium color, (3) visible turbidity or flocculent formation, and (4) changes in the color or morphology of oil droplets.

### Bacterial isolation and verification of JP-5 degradation capabilities

After 4 weeks of incubation with JP-5 jet fuel, samples were mixed with 40% glycerol and stored at –80°C for future analysis. To isolate individual bacterial colonies, 10 μL aliquot from JP-5 amended positive MPN tubes amended with JP-5 were spread onto Trypticase soy broth (TSB, Difco Laboratories, MI, USA) agar plates using the spread plate technique. 8-10 single colonies were randomly selected from 12 soil samples (total of 141 isolates). Each colony was purified through three successive rounds of streak plating on TSA agar to obtain pure bacterial isolates.

To evaluate their ability to grow with JP-5 as the sole carbon source, these purified isolates were inoculated into fresh BH medium supplemented with 1% (v/v) JP-5 jet fuel. Cultures were incubated at 30°C with shaking at 180 rpm for four weeks, and growth was monitored daily by measuring OD_600_ using a spectrophotometer (Bausch & Lomb, Laval, Canada). Two types of negative controls were included: (1) BH medium with 1% (v/v) JP-5 but no bacterial inoculum, and (2) BH medium inoculated with soil samples but without JP-5. Both negative controls consistently showed no detectable OD_600_ signal over the 4-week incubation period.

### Modeling of bacterial growth

Bacterial growth curves in the batch reactors with JP-5 as the sole carbon source were modeled using the logistic growth model: 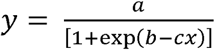 where *y* represents the OD_600_, and *x* denotes time. The model includes three parameters: *a*, the maximum OD_600_ value representing the maximum optical density; *b*, a constant related to the relative growth rate; and *c*, the relative lag phase duration time at which the population reaches half of the carrying capacity (i.e., the inflection point of the sigmoid curve) (Zwietering et al., 1990). Curve fitting and parameter estimation were performed using MATLAB R2025a (MathWorks, MA, USA).

### 16S rRNA gene sequencing

A total of 35 JP-5 degrading bacterial isolates were selected from the top 2-3 fast-growing isolates from each soil sample for genomic DNA extraction and 16S rRNA gene sequencing for phylogenetic analysis. DNA was extracted from single colonies using a simple NaOH lysis method (Saingam et al., 2020), and the 16S rRNA gene was amplified with universal primers 27F (5’ AGRGTTYGATYMTGGCTCAG 3’) and 1492R (5’ RGYTACCTTGTTACGACTT 3’). PCR was performed with GoTaq DNA polymerase (Promega, WI, USA) using an initial denaturation at 95°C for 2 min, followed by 35 cycles of 95°C for 30 s, 57°C for 30 s, and 72°C for 1 min. Amplicons were confirmed on a 1.5% agarose gel, purified with QIAquick kit (Qiagen, Hilden, Germany), and sequenced by Sanger sequencing using Applied Biosystems 3730XL DNA Analyzer (Thermo Fisher Scientific, MA, USA) at the Advanced Studies in Genomics, Proteomics and Bioinformatics (ASGPB) facility, University of Hawaii at Mānoa.

### Quantifying biosurfactant production

Biosurfactant production by bacterial isolates was screened using the drop-collapse assay (Varadavenkatesan & Murty, 2013). Sterile glass slides were uniformly coated with a layer of commercial motor oil (Super Tech SAE 5W-20; Warren Oil, NC, USA) and left at room temperature for 24 hours to allow surface equilibration. Subsequently, 10 μL of cell-free culture supernatant was pipetted onto the oil-coated surface and incubated for 1 hour. The diameter of the drop was then measured. Each assay was performed in triplicate, and a negative control (10 μL of deionized water) was included in all assays to confirm assay specificity.

### Data analysis

Sanger sequencing data were obtained in chromatogram format (.ab files), analyzed using SnapGene software (GSL Biotech, CA, USA), and forward and reverse reads were merged through multiple sequence alignment by CLUSTAL W (Thompson et al., 1994). Taxonomic identification was performed by querying the merged sequences against the National Center for Biotechnology Information (NCBI) nucleotide database using the nucleotide Basic Local Alignment Search Tool (BLASTn). The 16S rRNA sequences were first aligned with Multiple Sequence Alignment by CLUSTALW and a maximum-likelihood phylogenetic tree was constructed in PhyML v3.0 (Guindon et al., 2010) using the GTR+G+I model with 1,000 bootstrap replicates. Phylogenetic tree visualization was performed using iTOL v7 (Letunic & Bork, 2024) and reference 16S rRNA FASTA files were downloaded from the NCBI nucleotide database. For phylogenetic tree construction, 16S rRNA sequences of *Escherichia coli* strain NBRC 102203 (GenBank accession: NR_114042.1) and *Mycolicibacterium vanbaalenii* strain PYR-1 (GenBank accession: NR_029293.1) were included as distantly related outgroup references. The sequencing data are available in the NCBI GenBank submission number under SUB15779560 with accession numbers from PX557911 to PX557945.

Spearman’s rank correlation analysis (MATLAB 2025a) is conducted between the drop collapse diameter with growth dynamic model parameters, including maximum optical density, relative growth rate, and relative lag phase duration.

## Results

### Abundance of JP-5-degrading bacteria in Oahu soil

The abundance of JP-5-degrading bacteria in the soil samples ranged widely, from 8.1 × 10^1^ MPN/g to 5.1 × 10^4^ MPN/g, with a geometric mean of 1.9 × 10^4^ MPN/g (σ= 3.4 × 10^4^ MPN/g; **Figure 2b**). Since various factors can contribute to the large variations observed, including the total bacterial biomass in the samples, the relative abundance of JP-5-degrading bacteria was estimated by calculating the concentration ratio between JP-5-degrading bacteria and total bacteria (**Figure 2a**). The relative abundance of JP-5 bacteria which from 0.003% to 11.1%, with a geometric mean of 2.4% (σ= 4.4 %).

**Figure 2.**
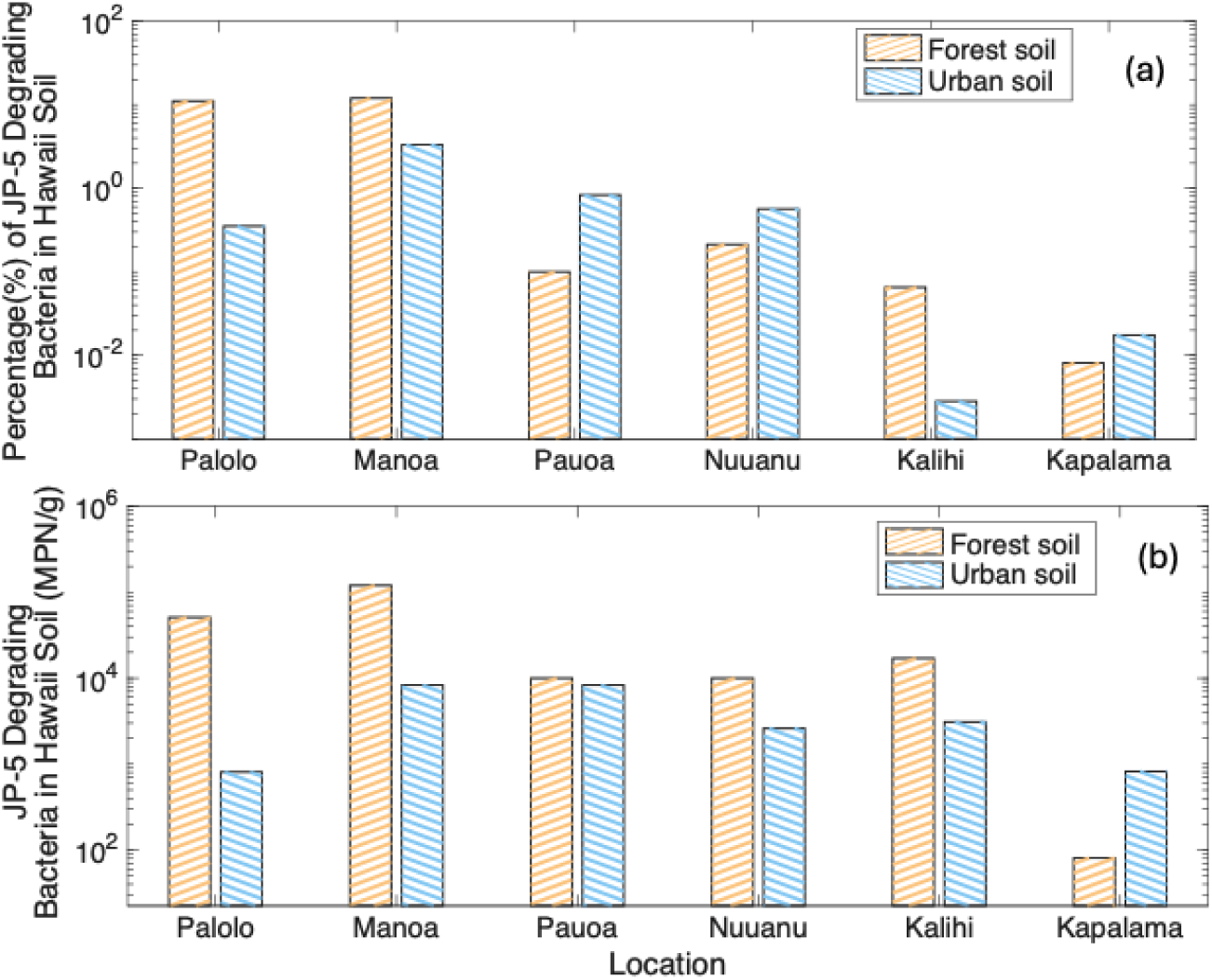
Relative abundance of JP-5 degrading bacteria in 12 locations (Forest soil and Urban soil).

### Phylogenetic diversity of JP-5-degrading bacteria

Most of the JP-5-degrading soil bacterial isolates were from *Achromobacter* (*n* = 13; 37%)*, Ralstonia* (*n* = 8; 23%)*, Pseudomonas* (*n* = 5; 14%), and *Stenotrophomonas* (*n* = 5; 14%) (**Figure 3**). There were also four bacterial isolates each belonging to *Acinetobacter* (*n* = 1)*, Staphylococcus* (*n* = 1)*, Bacillus* (*n* = 1), and *Gordonia* (*n* = 1). Among these genera, only a few isolates could be confidently assigned to the species level. Specifically, several *Pseudomonas* isolates were identified as *P. aeruginosa*, and one *Staphylococcus* isolate was identified as *S. aureus*. Within the *Achromobacter* and *Stenotrophomonas* isolates, multiple distinct 16S rRNA sequence types were observed, indicating intra-genus diversity. These results suggest that JP-5 degradation in these soils is primarily associated with a limited number of dominant genera, while several other taxa may play minor or supportive roles in the community. The distribution of isolates across genera also varied among the sampling sites, highlighting potential differences in microbial community composition and hydrocarbon-degrading potential.

**Figure 3.**
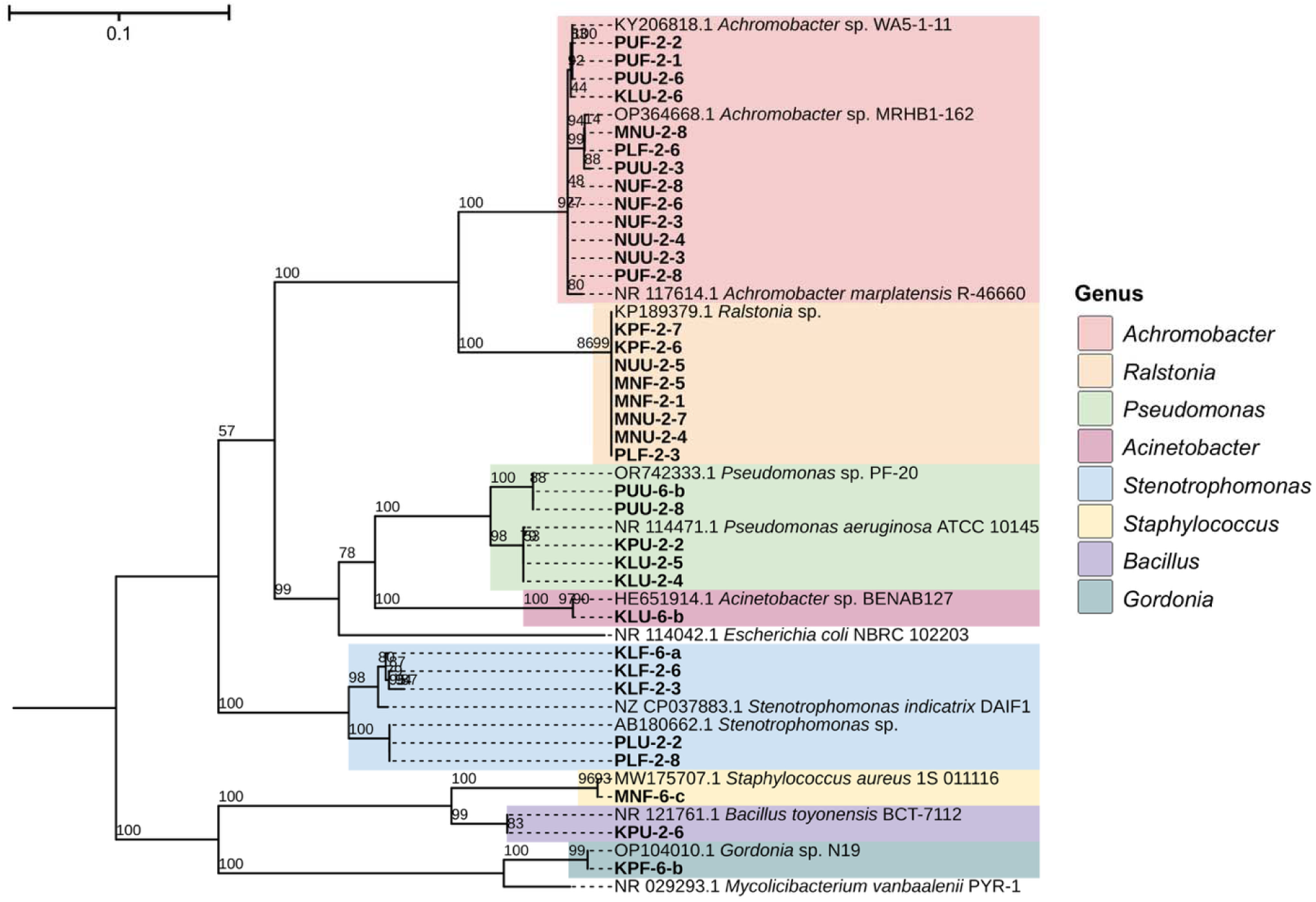
Maximum-likelihood phylogenetic tree based on the alignment of nearly full-length 16S rRNA gene sequences from 35 bacterial Reference sequences were retrieved from the NCBI nucleotide database to support taxonomic classification and assess relatedness. Isolate names are labeled accordingly, and taxa representing different genera are indicated with color-coded labels. The tree was rooted using *Escherichia coli* NBRC 102203 and *Mycolicibacterium vanbaalenii* PYR-1 as outgroup references.

### Bacterial growth kinetics with JP-5 as the sole carbon source

The logistic growth model was used to model the batch growth curves of JP-5-degrading bacteria when JP-5 was provided as the sole carbon source. Key bacterial growth parameters, including maximum optica density (a), relative growth rate (b), and relative lag phase duration (c), were determined (**Figure 4a).** The modeling was illustrated using representative growth curves from *Pseudomonas* isolate, demonstrated rapid growth within the first 3–4 days, followed by a plateau phase indicating stabilization of cell density after approximately one week (**Figure 4b**). In contrast, the *Gordonia* isolate, displayed a delayed growth pattern, with a marked increase in OD_600_ occurring between days 16–20 and reaching its maximum OD_600_ around day 22 (**Figure 4c**). The relative growth rate of *Pseudomonas* was 0.15 OD_600_ day ^-1^, compared to a slightly lower rate of 0.08 OD_600_ day^-1^ for *Gordonia*. The maximum optical density of *Pseudomonas* reached 0.22 OD_600_, whereas Gordonia exhibited a lower maximum optical density of 0.12 OD_600_.

**Figure 4.**
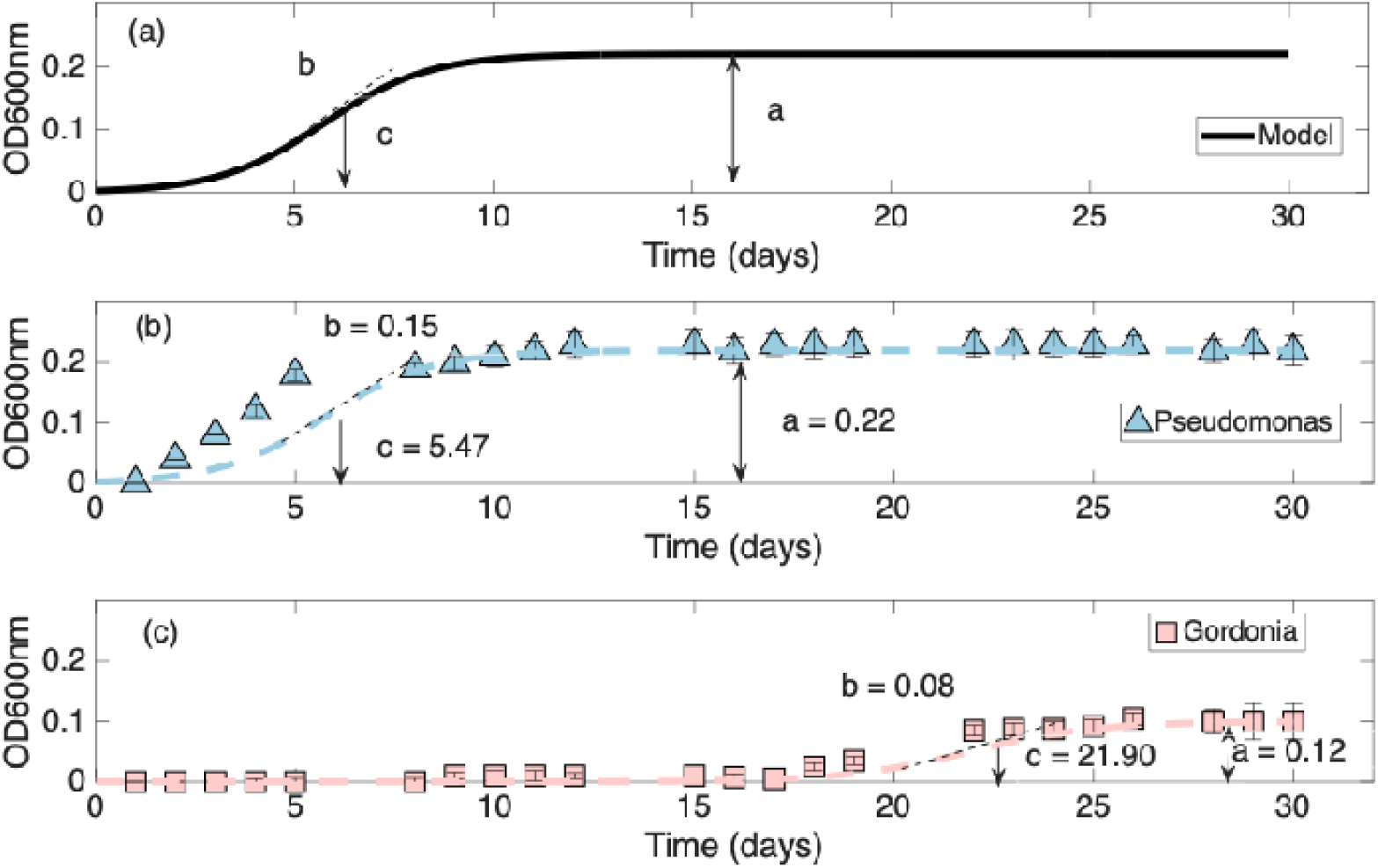
Modeling of example growth curve of the isolates from *Pseudomonas* sp. (blue line, filled triangle markers) and *Gordonia* sp. (pink line, square markers) with JP-5 as the sole carbon source.

The logistic growth model was used to fit the batch growth data of 39 JP-5-degrading bacterial isolates from Hawaii soil. The growth kinetic parameters and corresponding R^2^ for all isolates are provided in **Table S2**. **Figure 5** illustrates the distribution of growth parameters, maximum optical density (**Figure 5a**), relative growth rate (**Figure 5b**), and relative lag phase duration (**Figure 5c**) in relation to the phylogenetic affiliation of the JP-5-degradaing bacterial isolates. The maximum optical density ranged from 0.06 OD_600_ to 0.82 OD_600_ across isolates from all bacterial isolates, with a geometric mean of 0.18 ((J = 0.09). Relative growth ranging from 0.03 OD_600_ to 0.64 OD_600_ day ^1^with a geometric mean of 0.26 OD_600_ day^-1^ ((J = 0.19). The lag time ranges from 2.0 -24 .0 days with a geometric mean of 8.0 day ((J = 4.5).

**Figure 5.**
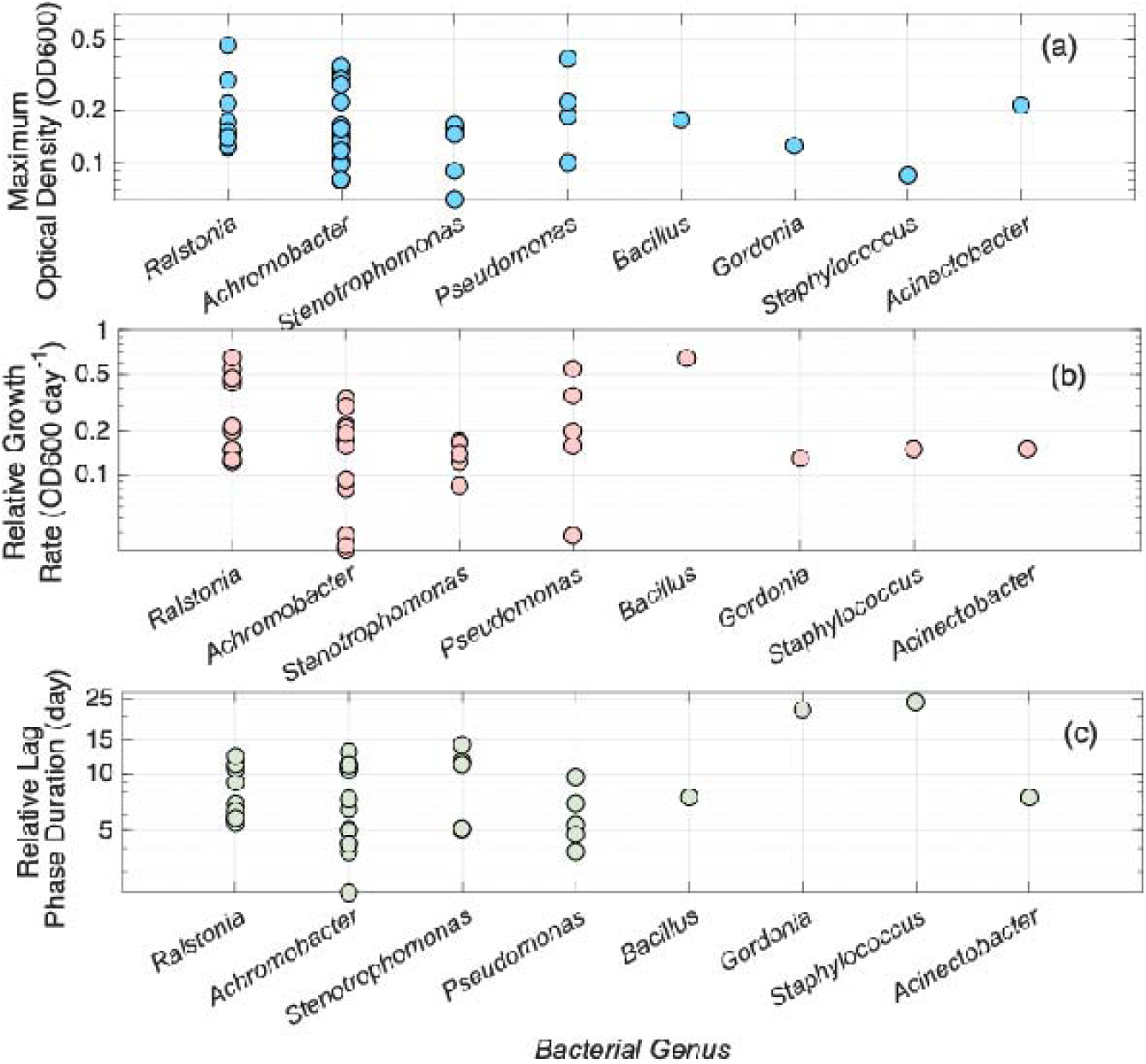
The average maximum optical density (a), relative growth rate (b), and relative lag phase duration (c) of the bacterial isolates were compared among bacterial genera.

### Bacterial biosurfactant production

Biosurfactant production by the JP-5-degrading bacterial isolates, which was quantified using the drop collapse assay (Varadavenkatesan & Murty, 2013) were analyzed with respect to the bacterial growth parameters, including maximum optical density, relative growth rate, and relative lag phase duration (**Figure 6**). The analysis revealed that higher biosurfactant production was generally associated with increased maximum optical density (**Figure 6a**, *r* = 0.69, *p* < 0.001) and relative growth rate (**Figure 6b**, *r* = 0.41, *p* = 0.008) but inversely correlated with relative lag phase duration (**Figure 6c**, *r* = –0.36, *p* = 0.02).

**Figure 6.**
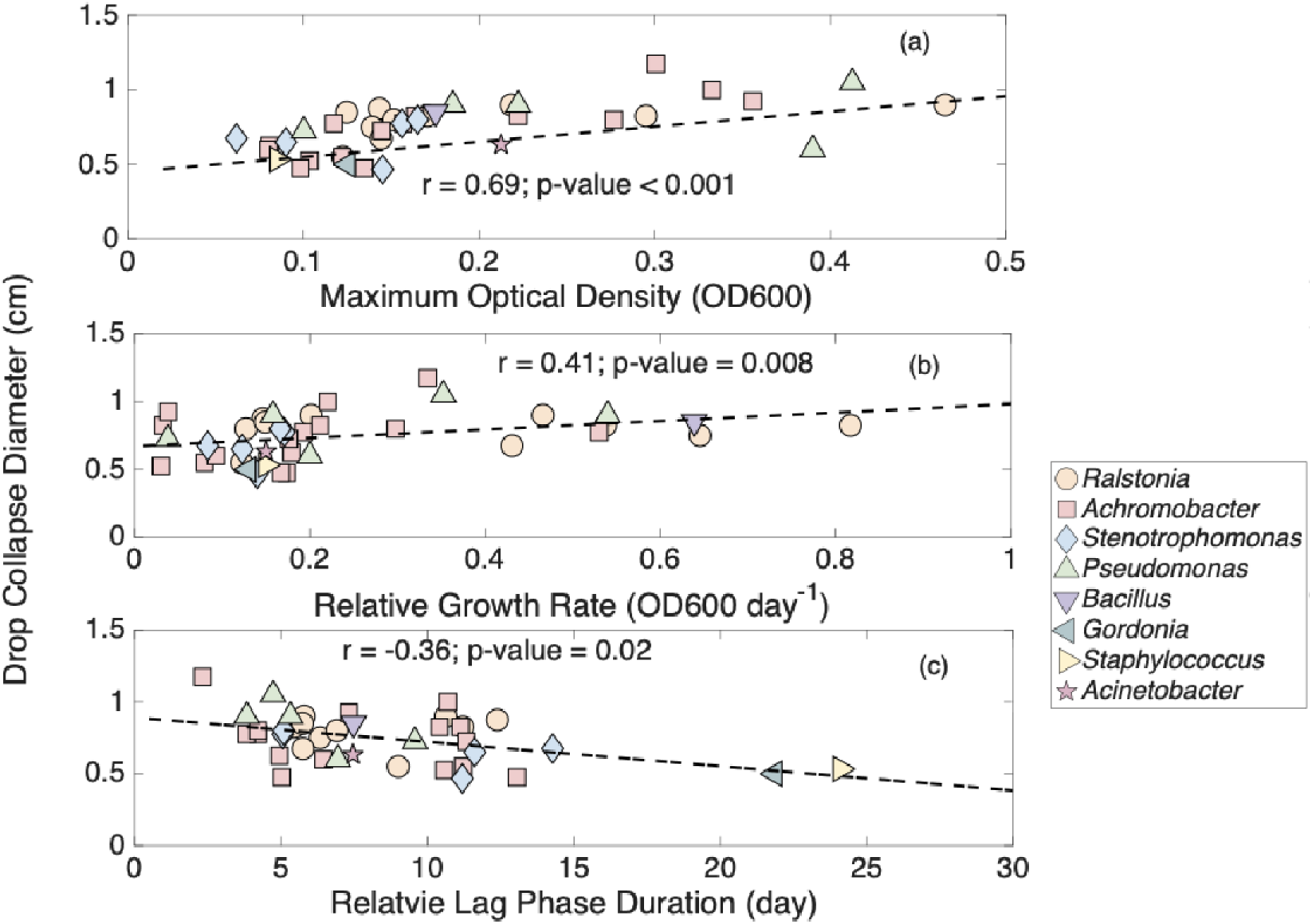
Relationship between the average of drop collapse diameter (cm) and maximum optical density (a), relative growth rate (b), and relative lag phase duration (c) as indicated by Spearman’s rank correlation coefficients. Shapes and colors represent different bacteria genera.

## Discussion

The spatial distribution of JP-5-degrading bacteria across the six O’ahu watersheds revealed difference between forest and urban soils (**Figure 2**). Palolo, Manoa and Kalihi displayed markedly elevated proportions of degraders, consistent with the positive correlation observed between degrader abundance and soil water content (**Figure S1**). In contrast, urban soils generally showed reduced percentage of JP-5 degraders at these locations, possibly due to lower moisture, or inhibitory contaminants associated with developed environments.

The genetic diversity observed among the isolated genera suggests potential functional variability and adaptation to distinct hydrocarbon substrates and soil microenvironments (**Figure 3**). This taxonomic diversity highlights a metabolically complex soil microbiome capable of degrading JP-5 PHCs. *Pseudomonas* is well-known bacterial genus capable of metabolizing *n*-alkanes, which are major constituents of jet fuel, and can produce biosurfactants that enhance hydrocarbon bioavailability (Gunasekera et al., 2013). *Stenotrophomonas*, although less frequently studied, has emerged as a metabolically versatile genus capable of degrading both aliphatic and aromatic hydrocarbons and is frequently detected in consortia inhabiting hydrocarbon-contaminated sites (Juhasz et al., 2000).

*Achromobacter* was the most frequently isolated genus in the Hawaii soil samples, which also exhibited significant diversity as indicated by the five distinct 16S rRNA sequence types among the isolates assigned to the *Achromobacter* genus. Previous studies have shown that the *Achromobacter* genus exhibits metabolic versatility, capable of degrading mid-chain diesel-range hydrocarbons (C_14_ – C_19_) (Bordoloi et al., 2014). *Ralstonia* was the second most common genus detected in the Hawaii soil, which has also been frequently reported to degrade crude oil and branched alkanes (Purnomo et al., 2019).

*Acinetobacter* species possess key enzymatic systems, such as alkane hydroxylases *alkB* and *almA*, enabling them to degrade *n*-alkanes (J Czarny et al., 2020). Members of the genus *Bacillus* contribute significantly to hydrocarbon degradation by producing potent biosurfactants that increase the solubility and bioavailability of jet fuel constituents (Parthipan et al., 2017).

*Gordonia* species, equipped with monooxygenases and cytochrome P450 enzymes, play an essential role in the oxidation of long-chain n-alkanes (up to C_36_) commonly present in aviation fuel (Lo Piccolo et al., 2011). Although *Staphylococcus aureus* is not generally regarded as a primary hydrocarbon degrader, some *Staphylococcus* species have been isolated from petroleum-contaminated environments and aviation fuel tanks, indicating their tolerance to hydrocarbons and limited capacity to metabolize lighter fractions such as alkanes or aromatic intermediates (Rauch et al., 2006). Collectively, the presence of these diverse bacterial taxa indicates the ecological and metabolic diversity of microbial populations capable of adapting to and degrading jet fuel hydrocarbons.

The growth dynamics of JP-5 degrading bacterial isolates varied substantially across bacterial genera, reflecting functional and physiological difference in how these taxa metabolize jet fuel (**Figure 5**). *Ralstonia*, *Achromobacter* and *Pseudomonas* displayed relatively high maximum optical density values (**Figure 5a**), indicating their ability to large amounts of biomass potentially due to efficient catabolic pathways when JP-5 serves as the sole carbon source. This aligns with prior studies describing these genera as efficient degraders of mid-chain alkanes and aromatic hydrocarbons (Medic et al., 2020; Purnomo et al., 2023; Wu et al., 2018). This was corroborated by the observed growth rate patterns (**Figure 5b**), where *Ralstonia*, *Achromobacter* and *Pseudomonas* displayed higher relative growth rates. Although *Gordonia* is known for its ability to degrade long-chain alkanes, the low growth rates observed may be due to persistent and slower degradation with long chain compounds or weaker capacity compared to classical fast degraders. Relative lag phase duration (**Figure 5c**) indicated that isolates with rapid growth tended to exhibit shorter lag times, while those with slower growth rate tend to have prolonged adaptation periods (e.g., *Gordonia* and *Staphylococcus*), it may indicate a more complex adaptation process (Liu et al., 2021). However, this pattern should be interpreted with caution, as it may reflect the restricted sample size of these bacterial genera.

The positive correlation between biosurfactant production with maximum optical density and relative growth rate (**Figure 6**) suggests that biosurfactant synthesis is likely responsible for enhanced hydrocarbon bioavailability. Previous studies have reported that biosurfactant synthesis is often growth-associated, occurring primarily during the exponential growth phase, where it serves as a secondary metabolite linked to cell division and nutrient assimilation (Ebadi et al., 2017; Ibrahim et al., 2020; Karlapudi et al., 2018; Li et al., 2022). By reducing surface and interfacial tension, biosurfactants enhance the emulsification of hydrophobic petroleum hydrocarbon compounds, thereby increasing their pseudo-solubility and bioavailability. This process facilitates more efficient substrate uptake and utilization, which can accelerate growth kinetics and reduce the duration of the lag phase (Tavares et al., 2024; Zargar et al., 2022).

### Limitations and future studies

One limitation of this study is the use of bacterial growth as a proxy for hydrocarbon degradation, rather than directly measuring the reduction of specific JP-5 components. JP-5 jet fuel is a complex mixture of hydrocarbons, including alkanes, cycloalkanes, and aromatics, each of which may degrade at different rates and via distinct metabolic pathways. Quantifying the degradation of individual components remains challenging but is essential for accurately assessing bioremediation potential. Future studies should incorporate analytical techniques such as gas chromatography-mass spectrometry (GC-MS) to directly measure the breakdown of representative JP-5 compounds. In addition to growth and biosurfactant production, the genetic mechanisms underlying hydrocarbon degradation were not examined. Whole-genome sequencing of key isolates would enable identification of functional genes involved in hydrocarbon metabolism (e.g., *alkB*, *cyp153*), providing insight into metabolic pathways and strain-specific capabilities.

Furthermore, future work should explore the formation and fate of intermediate transformation products that produce by different bacteria taxa during JP-5 biodegradation. Comparative analyses across key degraders (e.g, *Pseudomonas*, *Achromobacter*, *Ralstonia*) could help to elucidate species-specific metabolic pathways and further understand the biochemical mechanisms driving hydrocarbon breakdown. Finally, examining bacterial consortia, rather than individual strains, may better reflect natural conditions and reveal synergistic effects that enhance degradation efficiency.

## Conclusion

This study demonstrates the capacity of Hawaii soil microbiota to degrade JP-5 jet fuel, highlighting significant variability in the relative abundance of JP-5-degrading bacteria across 12 sampling sites. The relative abundance of JP-5-degrading bacteria was positively associated with soil water content, biochemical oxygen demand, and overall degrader abundance. Eight fast-growing bacterial genera were identified to use JP-5 as the sole carbon source: *Achromobacter*, *Ralstonia*, *Stenotrophomonas*, *Pseudomonas*, *Acinetobacter*, *Staphylococcus*, *Bacillus*, and *Gordonia*. Notably, distinct growth behaviors were observed between these bacterial isolates; *Achromobacter*, *Ralstonia* and *Pseudomonas* exhibited shorter lag phases and higher growth rates, whereas *Gordonia* and *Staphylococcus* displayed slower growth and extended adaptation periods, suggesting functional differentiation in hydrocarbon degradation strategies. Furthermore, the positive correlation between biosurfactant production and growth performance indicates that bacterial biosurfactants are involved in improved hydrocarbon bioavailability and subsequent microbial utilization. These findings indicate the ubiquitous presence and phylogenetic diversity of JP-5 degrading microbial capabilities in the Hawaii soil microbiome and underscore their potential for application in bioremediation.

## Acknowledgement

This work was funded by the U.S. Department of Defense, Office of Naval Research under grant number N00014-23-1-2738.

## Notes

### Competing Interest Statement

The authors have declared no competing interest.

## References

1. Araújo, W. J. D., Oliveira, J. S., Araújo, S. C. D. S., Minnicelli, C. F., Silva-Portela, R., Da Fonseca, M., Freitas, J. F., Silva-Barbalho, K., Napp, A. P., & Pereira, J. E. S. (2020). Microbial culture in minimal medium with oil favors enrichment of biosurfactant producing genes. Frontiers in Bioengineering and Biotechnology, 8, 962.

2. Bordoloi, N. K., Rai, S. K., Chaudhuri, M. K., & Mukherjee, A. K. (2014). Deep-desulfurization of dibenzothiophene and its derivatives present in diesel oil by a newly isolated bacterium Achromobacter sp. to reduce the environmental pollution from fossil fuel combustion. Fuel processing technology, 119, 236–244.

3. Brinkmann, M. T., Rong, K., Xie, Y., & Yan, T. (2024). Formation potential of disinfection byproducts during chlorination of petroleum hydrocarbon-contaminated drinking water. Chemosphere, 357, 142057. 10.1016/j.chemosphere.2024.142057

4. Bushnell, L., & Haas, H. (1941). The utilization of certain hydrocarbons by microorganisms. Journal of bacteriology, 41(5), 653–673.

5. Chunyan, X., Qaria, M. A., Qi, X., & Daochen, Z. (2023). The role of microorganisms in petroleum degradation: Current development and prospects. Sci Total Environ, 865, 161112. 10.1016/j.scitotenv.2022.161112

6. Czarny, J., Staninska-Pieta, J., Piotrowska-Cyplik, A., Juzwa, W., Wolniewicz, A., Marecik, R., Lawniczak, L., & Chrzanowski, L. (2020). Acinetobacter sp. as the key player in diesel oil degrading community exposed to PAHs and heavy metals. J Hazard Mater, 383, 121168. 10.1016/j.jhazmat.2019.121168

7. Diallo, M. M., Vural, C., Cay, H., & Ozdemir, G. (2021). Enhanced biodegradation of crude oil in soil by a developed bacterial consortium and indigenous plant growth promoting bacteria. J Appl Microbiol, 130(4), 1192–1207. 10.1111/jam.14848

8. Ebadi, A., Khoshkholgh Sima, N. A., Olamaee, M., Hashemi, M., & Ghorbani Nasrabadi, R. (2017). Effective bioremediation of a petroleum-polluted saline soil by a surfactant-producing Pseudomonas aeruginosa consortium. J Adv Res, 8(6), 627–633. 10.1016/j.jare.2017.06.008

9. Essaid, H. I., Bekins, B. A., Herkelrath, W. N., & Delin, G. N. (2011). Crude oil at the Bemidji site: 25 years of monitoring, modeling, and understanding. Groundwater, 49(5), 706–726. https://ngwa.onlinelibrary.wiley.com/doi/10.1111/j.1745-6584.2009.00654.x

10. Gibbs, R. A., & Hayes, C. R. (1988). The use of R2A medium and the spread plate method for the enumeration of heterotrophic bacteria in drinking water. Letters in Applied Microbiology, 6(2), 19–21. 10.1111/j.1472-765X.1988.tb01205.x

11. Guindon, S., Dufayard, J.-F., Lefort, V., Anisimova, M., Hordijk, W., & Gascuel, O. (2010). New algorithms and methods to estimate maximum-likelihood phylogenies: assessing the performance of PhyML 3.0. Systematic biology, 59(3), 307–321.

12. Gunasekera, T. S., Striebich, R. C., Mueller, S. S., Strobel, E. M., & Ruiz, O. N. (2013). Transcriptional profiling suggests that multiple metabolic adaptations are required for effective proliferation of Pseudomonas aeruginosa in jet fuel. Environmental science & technology, 47(23), 13449–13458.

13. Ibrahim, S., Abdul Khalil, K., Zahri, K. N. M., Gomez-Fuentes, C., Convey, P., Zulkharnain, A., Sabri, S., Alias, S. A., Gonzalez-Rocha, G., & Ahmad, S. A. (2020). Biosurfactant Production and Growth Kinetics Studies of the Waste Canola Oil-Degrading Bacterium Rhodococcus erythropolis AQ5-07 from Antarctica. Molecules, 25(17). 10.3390/molecules25173878

14. Juhasz, A. L., Stanley, G. A., & Britz, M. L. (2000). Microbial degradation and detoxification of high molecular weight polycyclic aromatic hydrocarbons by Stenotrophomonas maltophilia strain VUN 10,003. Letters in applied microbiology, 30(5), 396–401.

15. Jung, C. M., Broberg, C., Giuliani, J., Kirk, L. L., & Hanne, L. F. (2002). Characterization of JP-7 jet fuel degradation by the bacterium Nocardioides luteus strain BAFB. J Basic Microbiol, 42(2), 127–131. 10.1002/1521-4028(200205)42:2<127::AID-JOBM127>3.0.CO;2-C

16. Karlapudi, A. P., Venkateswarulu, T. C., Tammineedi, J., Kanumuri, L., Ravuru, B. K., Dirisala, V. R., & Kodali, V. P. (2018). Role of biosurfactants in bioremediation of oil pollution-a review. Petroleum, 4(3), 241–249. 10.1016/j.petlm.2018.03.007

17. Letunic, I., & Bork, P. (2024). Interactive Tree of Life (iTOL) v6: recent updates to the phylogenetic tree display and annotation tool. Nucleic acids research, 52(W1), W78–W82.

18. Li, C., Zhou, Z. X., Jia, X. Q., Chen, Y., Liu, J., & Wen, J. P. (2013). Biodegradation of crude oil by a newly isolated strain Rhodococcus sp. JZX-01. Appl Biochem Biotechnol, 171(7), 1715–1725. 10.1007/s12010-013-0451-4

19. Li, Z., Rosenzweig, R., Chen, F., Qin, J., Li, T., Han, J., Istvan, P., Diaz-Reck, D., Gelman, F., Arye, G., & Ronen, Z. (2022). Bioremediation of Petroleum-Contaminated Soils with Biosurfactant-Producing Degraders Isolated from the Native Desert Soils. Microorganisms, 10(11). 10.3390/microorganisms10112267

20. Liu, B., Ju, M., Liu, J., Wu, W., & Li, X. (2016). Isolation, identification, and crude oil degradation characteristics of a high-temperature, hydrocarbon-degrading strain. Mar Pollut Bull, 106(1-2), 301–307. 10.1016/j.marpolbul.2015.09.053

21. Liu, Y., Wu, J., Liu, Y., & Wu, X. (2021). Biological process of alkane degradation by Gordonia sihwaniensis. ACS omega, 7(1), 55–63.

22. Lo Piccolo, L., De Pasquale, C., Fodale, R., Puglia, A. M., & Quatrini, P. (2011). Involvement of an alkane hydroxylase system of Gordonia sp. strain SoCg in degradation of solid n-alkanes. Applied and environmental microbiology, 77(4), 1204–1213.

23. Medic, A., Ljesevic, M., Inui, H., Beskoski, V., Kojic, I., Stojanovic, K., & Karadzic, I. (2020). Efficient biodegradation of petroleum n-alkanes and polycyclic aromatic hydrocarbons by polyextremophilic Pseudomonas aeruginosa san ai with multidegradative capacity. RSC Adv, 10(24), 14060–14070. 10.1039/c9ra10371f

24. Meintanis, C., Chalkou, K. I., Kormas, K. A., & Karagouni, A. D. (2006). Biodegradation of crude oil by thermophilic bacteria isolated from a volcano island. Biodegradation, 17(2), 105–111. 10.1007/s10532-005-6495-6

25. Mohammed, S. A., Omar Zrary, T. J., & Hasan, A. H. (2023). Degradation of crude oil and the pure hydrocarbon fractions by indigenous soil microorganisms. Biologia, 78(12), 3637–3651. 10.1007/s11756-023-01513-4

26. Pandolfo, E., Barra Caracciolo, A., & Rolando, L. (2023). Recent Advances in Bacterial Degradation of Hydrocarbons. Water, 15(2). 10.3390/w15020375

27. Papen, H., & Von Berg, R. (1998). A most probable number method (MPN) for the estimation of cell numbers of heterotrophic nitrifying bacteria in soil. Plant and soil, 199(1), 123–130.

28. Parthipan, P., Preetham, E., Machuca, L. L., Rahman, P. K., Murugan, K., & Rajasekar, A. (2017). Biosurfactant and degradative enzymes mediated crude oil degradation by bacterium Bacillus subtilis A1. Frontiers in Microbiology, 8, 193.

29. Posada-Baquero, R., Jimenez-Volkerink, S. N., Garcia, J. L., Vila, J., Cantos, M., Grifoll, M., & Ortega-Calvo, J. J. (2020). Rhizosphere-enhanced biosurfactant action on slowly desorbing PAHs in contaminated soil. Sci Total Environ, 720, 137608. 10.1016/j.scitotenv.2020.137608

30. Purnomo, A. S., Putra, S. R., Putro, H. S., Hamzah, A., Rohma, N. A., Rohmah, A. A., Rizqi, H. D., Asranudin, Tangahu, B. V., Warmadewanthi, I., & Shimizu, K. (2023). The application of biosurfactant-producing bacteria immobilized in PVA/SA/bentonite bio-composite for hydrocarbon-contaminated soil bioremediation. RSC Adv, 13(31), 21163–21170. 10.1039/d3ra02249h

31. Purnomo, A. S., Rizqi, H. D., Harmelia, L., Anggraeni, S. D., Melati, R. E., Damayanti, Z. H., Shafwah, O. M. A., & Kusuma, F. C. (2019). Biodegradation of crude oil by Ralstonia pickettii under high salinity medium. Malays. J. Fundam. Appl. Sci, 15, 377–380.

32. Rauch, M. E., Graef, H. W., Rozenzhak, S. M., Jones, S. E., Bleckmann, C. A., Kruger, R. L., Naik, R. R., & Stone, M. O. (2006). Characterization of microbial contamination in United States Air Force aviation fuel tanks. Journal of Industrial Microbiology and Biotechnology, 33(1), 29–36.

33. Reasoner, D. J., & Geldreich, E. (1985). A new medium for the enumeration and subculture of bacteria from potable water. Applied and environmental microbiology, 49(1), 1–7.

34. Saingam, P., Li, B., & Yan, T. (2020). Fecal indicator bacteria, direct pathogen detection, and microbial community analysis provide different microbiological water quality assessment of a tropical urban marine estuary. Water research, 185, 116280.

35. Smith, K. W., Proctor, S. P., Ozonoff, A., & McClean, M. D. (2010). Inhalation exposure to jet fuel (JP8) among U.S. Air Force personnel. J Occup Environ Hyg, 7(10), 563–572. 10.1080/15459624.2010.503755

36. Song, X., Xu, Y., Li, G., Zhang, Y., Huang, T., & Hu, Z. (2011). Isolation, characterization of Rhodococcus sp. P14 capable of degrading high-molecular-weight polycyclic aromatic hydrocarbons and aliphatic hydrocarbons. Mar Pollut Bull, 62(10), 2122–2128. 10.1016/j.marpolbul.2011.07.013

37. Tavares, J., Paixao, S. M., Silva, T. P., & Alves, L. (2024). New Insights on Gordonia alkanivorans Strain 1B Surface-Active Biomolecules: Gordofactin Properties. Molecules, 30(1). 10.3390/molecules30010001

38. Thompson, J. D., Higgins, D. G., & Gibson, T. J. (1994). CLUSTAL W: improving the sensitivity of progressive multiple sequence alignment through sequence weighting, position-specific gap penalties and weight matrix choice. Nucleic acids research, 22(22), 4673–4680.

39. Van Hamme, J. D., Singh, A., & Ward, O. P. (2003). Recent advances in petroleum microbiology. Microbiol Mol Biol Rev, 67(4), 503–549. 10.1128/MMBR.67.4.503-549.2003

40. Varadavenkatesan, T., & Murty, V. R. (2013). Production of a Lipopeptide Biosurfactant by a Novel Bacillus sp. and Its Applicability to Enhanced Oil Recovery. ISRN Microbiol, 2013, 621519. 10.1155/2013/621519

41. Varjani, S. J., & Upasani, V. N. (2016). Biodegradation of petroleum hydrocarbons by oleophilic strain of Pseudomonas aeruginosa NCIM 5514. Bioresour Technol, 222, 195–201. 10.1016/j.biortech.2016.10.006

42. Wang, D., Lin, J., Lin, J., Wang, W., & Li, S. (2019). Biodegradation of Petroleum Hydrocarbons by Bacillus subtilis BL-27, a Strain with Weak Hydrophobicity. Molecules, 24(17). 10.3390/molecules24173021

43. Wu, T., Xu, J., Xie, W., Yao, Z., Yang, H., Sun, C., & Li, X. (2018). Pseudomonas aeruginosa L10: A Hydrocarbon-Degrading, Biosurfactant-Producing, and Plant-Growth-Promoting Endophytic Bacterium Isolated From a Reed (Phragmites australis). Front Microbiol, 9, 1087. 10.3389/fmicb.2018.01087

44. Zargar, A. N., Mishra, S., Kumar, M., & Srivastava, P. (2022). Isolation and chemical characterization of the biosurfactant produced by Gordonia sp. IITR100. PLoS One, 17(4), e0264202. 10.1371/journal.pone.0264202

45. Zwietering, M. H., Jongenburger, I., Rombouts, F. M., & Van’t Riet, K. (1990). Modeling of the bacterial growth curve. Applied and environmental microbiology, 56(6), 1875–1881.

