## Supplemental materials for "Characterizing JP-5 Biodegradation Potential in Hawaii Soil Microbiome: Phylogeny, Growth Kinetics, and Biosurfactant Production"

**Supplemental Data**

Table S1: Sampling locations, basic parameters, abundance and relative abundance of JP-5 degrading bacteria from the soil sample

| Locations  (Coordinate) | ID | pH | Water content (%) | BOD  (mg/L) | Total bacteria count (MPN/g) | Total JP-5 degrading bacteria count (MPN/g) | Relative abundance of JP-5 degrading bacteria (%) |
| --- | --- | --- | --- | --- | --- | --- | --- |
| Palolo_forest  (21.314031, -157.785942) | PLF | 7.38 | 56.75 | 2.36 | 4.6x10^5^ | 5.1x10^4^ | 11.08 |
| Palolo_urban  (21.299130, -157.795029) | PLU | 7.42 | 18.09 | 2.19 | 2.3x10^5^ | 8.1x10^2^ | 0.35 |
| Manoa_urban  (21.307946, -157.809109) | MNU | 7.37 | 25.56 | 2.2 | 2.5x10^5^ | 8.3x10^3^ | 3.32 |
| Manoa_forest  (21.334757, -157.800426) | MNF | 7.12 | 66.01 | 2.42 | 1.0x10^6^ | 1.2x10^5^ | 12 |
| Pauoa_urban  (21.318484, -157.849714) | PUU | 7.21 | 14.85 | 2.23 | 1.0x10^6^ | 8.3x10^3^ | 0.83 |
| Pauoa_forest  (21.327898, -157.832013) | PUF | 7.17 | 60.13 | 2.48 | 1.0x10^7^ | 1.0x10^4^ | 0.1 |
| Nuuanu_urban  (21.319560, -157.855624 ) | NUU | 7.30 | 27.74 | 1.69 | 4.6x10^5^ | 2.6x10^3^ | 0.57 |
| Nuuanu_forest  (21.346995, -157.821050) | NUF | 7.18 | 58.40 | 2.05 | 4.7x10^6^ | 1.0x10^4^ | 0.21 |
| Kalihi_urban  (21.342693, -157.868954) | KLU | 7.27 | 15.20 | 2.24 | 1.1x10^8^ | 3.1x10^3^ | 0.003 |
| Kalihi_forest  (21.364745, -157.841556) | KLF | 7.15 | 17.23 | 2.19 | 2.6x10^7^ | 1.7x10^4^ | 0.07 |
| Kapalama_urban  (21.329776, -157.866673) | KPU | 7.33 | 11.22 | 2.10 | 4.7x10^6^ | 8.2x10^2^ | 0.02 |
| Kapalama_forest  (21.338678, -157.855443) | KPF | 7.30 | 27.48 | 2.15 | 1.0x10^6^ | 8.1x10^1^ | 0.008 |

Soil pH was measured in accordance with EPA Method 9045D. Gravimetric water content was determined by drying 10 g of soil on aluminum trays at 105°C for 24 h and calculating the mass loss. Soil biological oxygen demand (BOD) was assessed by following the APHA Standard Method 9221; briefly, 3 g of soil samples were mixed with 300 mL of deionized water in BOD bottles, which were then incubated in the dark at 20°C for 5 days and dissolved oxygen (DO) changes were measured using a DO meter (Thermo Fisher Scientific, MA, USA). This indicate relative microbial oxygen consumption potential in soil.

Table S2: Growth model parameters, R^2^ and Average drop collapse diameter from the isolates

| Isolate ID | Average carrying capacity (a) | Average growth rate (b) | Average lag time (c) | R^2^ | Average Drop Collapse Diameter (cm) |
| --- | --- | --- | --- | --- | --- |
| PLF-2-3 | 0.12 | 0.12 | 9.00 | 0.98 | 0.55 |
| PLF-2-6 | 0.33 | 0.22 | 10.70 | 0.32 | 1.00 |
| PLF-2-8 | 0.09 | 0.12 | 11.61 | 0.78 | 0.65 |
| PLU-2-2 | 0.06 | 0.08 | 14.28 | 0.97 | 0.68 |
| MNU-2-4 | 0.14 | 0.43 | 5.75 | 0.98 | 0.68 |
| MUN-2-7 | 0.22 | 0.20 | 10.62 | 0.91 | 0.90 |
| MUN-2-8 | 0.22 | 0.21 | 10.42 | 0.98 | 0.83 |
| MNF-2-1 | 0.17 | 0.81 | 11.21 | 0.92 | 0.83 |
| MNF-2-3 | 0.12 | 0.08 | 11.22 | 0.98 | 0.55 |
| MNF-2-5 | 0.14 | 0.14 | 12.39 | 0.93 | 0.88 |
| PUU-2-3 | 0.14 | 0.18 | 11.33 | 0.96 | 0.73 |
| PUU-2-6 | 0.08 | 0.09 | 6.45 | 0.97 | 0.60 |
| PUU-2-8 | 0.10 | 0.04 | 9.57 | 0.96 | 0.73 |
| PUF-2-1 | 0.10 | 0.03 | 10.57 | 0.97 | 0.53 |
| PUF-2-2 | 0.10 | 0.17 | 5.03 | 0.97 | 0.48 |
| PUF-2-8 | 0.13 | 0.17 | 13.06 | 0.96 | 0.48 |
| NUU-2-3 | 0.36 | 0.04 | 7.32 | 0.86 | 0.93 |
| NUU-2-4 | 0.08 | 0.18 | 4.95 | 0.93 | 0.63 |
| NUU-2-5 | 0.12 | 0.15 | 5.73 | 0.95 | 0.85 |
| NUF-2-3 | 0.30 | 0.33 | 2.32 | 0.87 | 1.18 |
| NUF-2-6 | 0.16 | 0.03 | 11.13 | 0.99 | 0.83 |
| NUF-2-8 | 0.28 | 0.30 | 4.21 | 0.92 | 0.80 |
| KLU-2-4 | 0.18 | 0.54 | 3.83 | 0.98 | 0.90 |
| KUL-2-5 | 0.22 | 0.15 | 5.33 | 0.96 | 0.90 |
| KLU-2-6 | 0.16 | 0.15 | 3.82 | 0.89 | 0.78 |
| KLF-2-3 | 0.16 | 0.17 | 5.06 | 0.93 | 0.78 |
| KLF-2-6 | 0.16 | 0.17 | 5.09 | 0.88 | 0.80 |
| KLF-2-8 | 0.12 | 0.19 | 4.22 | 0.94 | 0.78 |
| KPU-2-2 | 0.41 | 0.35 | 4.73 | 0.93 | 1.05 |
| KPU-2-6 | 0.18 | 0.63 | 7.48 | 0.84 | 0.85 |
| KPF-2-7 | 0.15 | 0.13 | 6.91 | 0.71 | 0.80 |
| KPF-2-6 | 0.30 | 0.53 | 5.44 | 0.88 | 0.83 |
| KPF-2-7 | 0.14 | 0.64 | 6.34 | 0.84 | 0.75 |
| KPF-2-8 | 0.47 | 0.46 | 5.79 | 0.85 | 0.90 |
| MNF-6-a | 0.15 | 0.14 | 11.19 | 0.97 | 0.47 |
| PUU-6-b | 0.13 | 0.13 | 21.85 | 0.32 | 0.50 |
| KLU-6-c | 0.09 | 0.15 | 24.11 | 0.77 | 0.53 |
| KLF-6-b | 0.39 | 0.20 | 6.94 | 0.97 | 0.60 |
| KPF-6-b | 0.21 | 0.15 | 7.46 | 0.98 | 0.63 |

Table S3: 16S rRAN result from fast growing and most abundant group (see excel file)

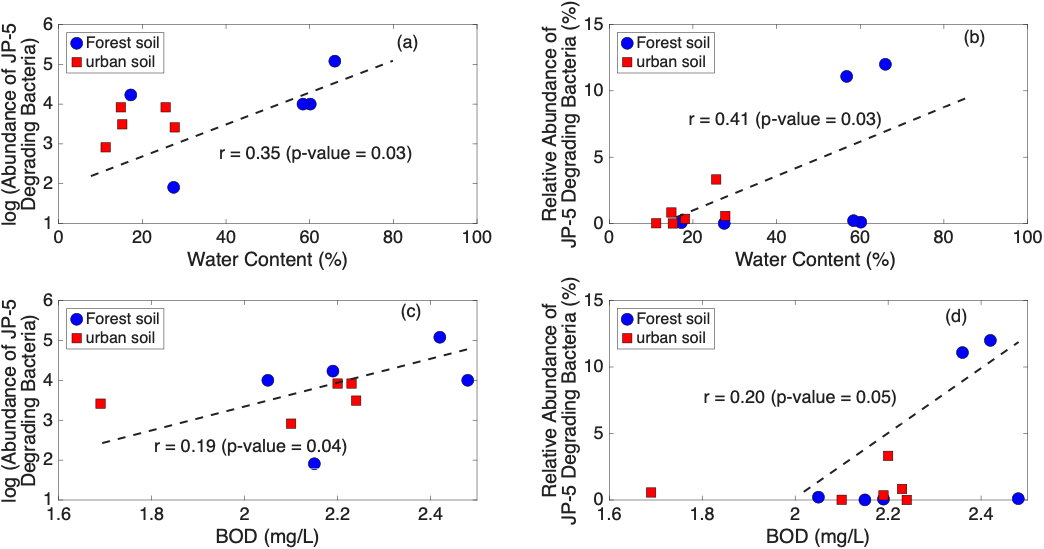

**Figure. S1** Abundance (left y-axis) and relative abundance (right y-axis) of JP-5 degrading bacteria show in relation to (a) water content (%) and (b) BOD

The large variations in the absolute abundance and the relative abundance of JP-5-degrading bacteria in the soil samples were analyzed against soil physicochemical parameters. Among the soil samples from the twelve sites, soil pH ranged from 7.1 to 7.4, while moisture content varied widely from 11.2% to 66.0%, with forest soils exhibiting higher water content than urban soils. BOD values ranged from 1.7 to 2.5 mg/L. The total and relative abundances of JP-5-degrading bacteria showed significant positive correlations with soil moisture (*r* = 0.35, *p* = 0.03, **Figure. S1a**; *r* = 0.41, *p* = 0.03, **Figure. S1b**). A similar trend was observed with BOD, where both total abundance and relative abundance of degraders were positively associated (*r* = 0.19, *p* = 0.04, **Figure S1c;** *r* = 0.20, *p* = 0.05, **Figure. S1d**).

The significant variations in the biomass of JP-5 degrading bacteria in the Oahu soil samples are likely the result of variations in key environmental factors that control natural soil bacterial biomass. This was supported by the significant positive correlation between the abundance of JP-5 degrading bacterial biomass and soil moisture content (**Figure. S1a**), as soil moisture is a well-known factor enhances hydrocarbon degradation efficiency (Mekonnen et al., 2024; Southwell et al., 2023). This was further supported the higher correlation coefficient between the relative abundance of JP-5 degrading biomass and soil water content (**Figure. S2b**). In comparison, the BOD content of the soil samples exhibited significant yet smaller correlation coefficients with the abundance of JP-5 degrading bacterial (**Figure. S1c&d**), indicating ubiquitous presence of JP-5-degrading bacteria in the soil microbiome.
